# Selective targeting of α_4_β_7_/MAdCAM-1 axis suppresses liver fibrosis by reducing proinflammatory T cell recruitment to the liver

**DOI:** 10.1101/2023.02.20.528201

**Authors:** Biki Gupta, Ravi Prakash Rai, Pabitra B. Pal, Sudrishti Chaudhary, Anna Chiaro, Shannon Seaman, Aatur D. Singhi, Satdarshan P. Monga, Smita S. Iyer, Reben Raeman

## Abstract

Integrin α_4_β_7_+ T cells perpetuate tissue injury in chronic inflammatory diseases, yet their role in hepatic fibrosis progression remains poorly understood. Here we report increased accumulation of α_4_β_7_+ T cells in the liver of people with cirrhosis relative to disease controls. Similarly, hepatic fibrosis in the established mouse model of CCl_4_-induced liver fibrosis was associated with enrichment of intrahepatic α_4_β_7_+ CD4 and CD8 T cells. Monoclonal antibody (mAb)-mediated blockade of α_4_β_7_ or its ligand mucosal addressin cell adhesion molecule (MAdCAM)-1 attenuated hepatic inflammation and prevented fibrosis progression in CCl_4_ treated mice. Improvement in liver fibrosis was associated with a significant decrease in the infiltration of α_4_β_7_+ CD4 and CD8 T cells suggesting that α_4_β_7_/MAdCAM-1 axis regulates both CD4 and CD8 T cell recruitment to the fibrotic liver, and α_4_β_7_+ T cells promote hepatic fibrosis progression. Analysis of hepatic α_4_β_7_+ and α_4_β_7_-CD4 T cells revealed that α_4_β_7_+ CD4 T cells enriched for markers of activation and proliferation demonstrating an effector phenotype. Notably, blockade of α_4_β_7_ or MAdCAM-1 did not affect the recruitment of Foxp3+ regulatory T cells, demonstrating the specificity of α_4_β_7_/MAdCAM-1 axis in regulating effector T cell recruitment to the liver. The findings suggest that α_4_β_7_+ T cells play a critical role in promoting hepatic fibrosis progression, and mAb-mediated blockade of α_4_β_7_ or MAdCAM-1 represents a promising therapeutic strategy for slowing hepatic fibrosis progression in chronic liver diseases.

## INTRODUCTION

Wound healing and repair are fundamental biological processes critical to maintaining tissue architecture and restoring organ homeostasis following injury ^1^. Protracted injury, however, can dysregulate this process resulting instead in excessive extracellular matrix deposition and tissue scarring or fibrosis. Progressive fibrosis is characteristic of advanced chronic liver diseases (CLDs), including alcoholic and nonalcoholic steatohepatitis as well as viral and autoimmune hepatitis^2–4^. Chronic unresolved fibrosis can lead to cirrhosis, the 11^th^ leading cause of global deaths, and a major risk factor for developing hepatocellular carcinoma (HCC) which is among the top 20 causes of deaths worldwide. Consequently, therapeutic strategies to stop progression of fibrosis are urgently needed.

The inflammatory response to liver injury, essential for resolution of injury and tissue repair, is a highly regulated process where both the initiation and resolution of inflammation are mediated by the liver-resident and liver-recruited immune cells. Chronic injury, however, leads to a perpetuation of hepatic inflammation characterized by sustained infiltration of immune cells to the liver and maintenance of fibrogenic pathways. While the liver-resident macrophages or Kupffer cells (KC) play a key role in the initiation of inflammation, recent data show that they are depleted in the advanced stages of CLD. Ultimately, the recruited proinflammatory immune cells including cells of both myeloid and lymphoid origin play a more prominent role in fibrosis progression making them an interesting target for therapeutic approaches to address fibrosis progression ^5^.

Immune cell recruitment to the liver following injury is mediated by chemokines and cytokines released by damaged hepatocytes, activated KCs as well as HSC-derived myofibroblasts. Recently, we demonstrated that the heterodimeric integrin receptor α_4_β_7_expressed on T cells and its ligand, mucosal addressin cell adhesion molecule (MAdCAM)-1, expressed on endothelial cells, drive hepatic inflammation in NASH by promoting recruitment of α_4_β_7_+ CD4 T cells to the liver ^6^. The α_4_β_7_/MAdCAM-1 axis is also implicated in promoting hepatic inflammation in chronic inflammatory liver diseases and primary sclerosing cholangitis (PSC), and upregulation of MAdCAM-1 expression is reported in the liver of most CLD patients ^7–10^. Despite strong evidence suggesting that the α_4_β_7_/MAdCAM-1 axis is influential in CLD, role of α_4_β_7_/MAdCAM-1 axis in promoting hepatic fibrosis progression remain poorly delineated.

In the current study, using established model of CCl_4_-induced liver fibrosis, we show a critical role of α_4_β_7_+CD4 and α_4_β_7_+CD8 T cells in promoting hepatic fibrosis progression. Our findings suggests that blocking α_4_β_7_/MAdCAM-1-mediated recruitment of α_4_β_7_+ T cells to the liver may represent a novel therapeutic strategy to slow or prevent fibrosis progression in CLD.

## METHODS

### Mice

Adult male C57BL/6J mice were obtained from The Jackson Laboratory (Bar Harbor, ME). The mice were maintained at the University of Pittsburgh Division of Animal Resources. Age- and sex-matched littermates were used for all experiments at 8–10 wk of age. All animal studies were carried out in accordance with protocols approved by the Institutional Animal Care and Use Committee at the University of Pittsburgh.

### Mouse model of liver fibrosis and α4β7 and MAdCAM-1 antibody treatment

To induce hepatic fibrosis, eight-week-old male mice were given twice-weekly oral gavage of CCl_4_ (2 mL/Kg CCl_4_ in 1:1 v/v olive oil) for six weeks. Mice receiving equal volume of olive oil served as vehicle controls. After 2 weeks of CCl_4_ treatment, a cohort of mice were treated with twice weekly intraperitoneal injections of 8 mg/Kg α_4_β_7_ mAb (DATK32; Bioxcell, West Lebanon, NH) or MAdCAM-1 mAb (MECA-367; Bioxcell, West Lebanon, NH) or rat IgG2a isotype antibody (Bioxcell, West Lebanon, NH) for four weeks.

### Human tissue

De-identified, fixed explanted liver tissues from people receiving orthotopic liver transplantation for decompensated liver cirrhosis were obtained from Biospecimen Processing and Repository Core at Pittsburgh Liver Research Centre. Liver tissue from tumor-adjacent parenchyma of people with hepatocellular carcinoma was used as controls.

### Histopathology

Histopathological analyses, including Hematoxylin and eosin (H&E) and Sirius Red staining, were conducted on formalin-fixed liver tissue sections as previously reported.^11^ Photomicrographs of the histologic sections were obtained by using a Zeiss Light Microscope (Zeiss, Jena, Germany).

### Immunofluorescence microscopy

Immunofluorescence (IF) microscopy was conducted on liver cryosections as previously reported.^11^ The IF images were visualized and obtained by using an Axioskop 2 plus microscope (Zeiss, Jena, Germany). Following antibodies were used for mouse tissue, α_4_β_7_/LPAM1(cat. No. 120702, Clone: MECA-367, Biolegend), aSMA (cat. No. MA1-06110, ThermoFisher Scientific), MAdCAM1(cat. No. 16-5997-85, ThermoFisher Scientific). Following antibodies were used for human tissue, α_4_β_7_/LPAM1(cat. No. DDX1434P-100, Clone: 11D9.03, Novus Biologicals) and CD3 (cat. No. <u>ab135372, Abcam</u>).

### Serological analysis

Serum alanine aminotransferase (ALT), and aspartate aminotransferase (AST) concentrations were measured using an AST and ALT Activity Assay Kit (Sigma-Aldrich, St. Louis, MO).

### Flow cytometric analysis

Livers perfused with 1X PBS (Thermo Fisher Scientific, Waltham, MA) were digested with 2mg/mL type IV collagenase (Worthington, NY) to obtain single cell suspensions and the lymphocytes were enriched by Percoll gradient centrifugation as describes previously ^11^. Enriched lymphocytes were stained with fluorochrome-conjugated antibodies and fixable viability stain (BD Biosciences, San Jose, CA), and samples were acquired on Cytek Aurora Spectral Cytometer (Cytek Biosciences, Bethesda, MD) equipped with five lasers. Data were analyzed using FlowJo software v X.10 (Tree Star, Inc., Ashland, OR) as describes previously ^11^. Total numbers were computed by multiplying proportion of specific populations to total numbers of enriched lymphocytes/liver. Fuorochrome conjugated antibodies against CD45 (30-F11), CD3 (17A2), CD4 (RM4-5), CD8 (53-6.7), CD44 (IM7), CD69 (H1.2F3), Tbet (4B10) and α_4_β_7_ (LPAM-1; DATK32) were purchased from BD Biosciences (San Jose, CA), and Foxp3 (FJK-16s), Ki67 (SolA15) and Live/Dead Aqua was purchased form ThermoFisher Scientific (Waltham, MA).

### Quantitative real-time PCR

Isolation of total RNA from liver, cDNA synthesis and qRT-PCR were performed as previously described.^11^ Data were normalized to 18S rRNA and data are presented as fold change in gene expression compared to controls.

### Statistical analysis

Two-tailed Student’s t test was employed to compare mean differences between two groups. One-way ANOVA in conjunction with post-hoc analysis for multiple group comparison was employed to investigate statistical differences. A *p* value < 0.05 was considered statistically significant. Data shown are representative of 3 independent experiments. Statistical analyses were performed using GraphPad Prism 8.0 (GraphPad Software) software.

## RESULTS

### Integrin α_4_β_7_ blockade attenuates CCl_4_-induced hepatic inflammation

We recently demonstrated that blocking α_4_β_7_+ T cell recruitment to the liver reduces hepatic injury in a murine model of NASH ^6^. To determine whether α_4_β_7_+ T cells play a role in promoting hepatic fibrosis, we used established mouse model of liver fibrosis to investigate whether blocking recruitment of α_4_β_7_+ T cells would be effective in decreasing hepatic fibrosis progression (**Fig. 1A**). Mice were treated with CCl_4_ for six weeks to induce hepatic fibrosis. Mice administered the vehicle olive oil served as controls. A cohort of CCl_4_-treated mice received α_4_β_7_ mAb for four weeks starting at week three. Mice treated with IgG isotype antibody served as controls (CCl_4_ + IgG). As shown in **Fig. 1B**, CCl_4_ + IgG-treated mice experienced severe liver injury indicated by increased infiltration of immune cells, steatosis and necrotic hepatocytes relative to vehicle controls. Increased liver injury in CCl_4_ + IgG-treated mice was further corroborated by high serum aspartate aminotransferase (AST) and alanine aminotransferase (ALT) levels relative to vehicle controls (**Fig. 1C-D**). Transcript levels of key pro-inflammatory cytokines, tumor necrosis factor-α (TNFα) and interleukin IL-6, were also significantly upregulated in the livers of CCl_4_ + IgG-treated mice relative to controls indicative of severe hepatic inflammation (**Fig. 1E-F**). In contrast, histological analysis of liver tissue sections from CCl_4_ + α_4_β_7_-mAb-treated mice revealed substantial improvement in hepatic inflammation as evidenced by reduced infiltration of immune cells and reduced tissue injury (**Fig. 1B**). The marked attenuation of liver injury in CCl_4_ + α_4_β_7_-mAb treated mice was further substantiated by significantly reduced serum ALT and AST levels (**Fig. 1C-D**). Compared to IgG controls, α_4_β_7_mAb treatment also significantly reduced transcript levels of key pro-inflammatory cytokines, TNF-α and IL-6, in the liver (**Fig. 1E-F**). Collectively, these results demonstrate that α_4_β_7_ blockade protected mice from CCl_4_-induced hepatic inflammation and injury.

**Figure 1.**
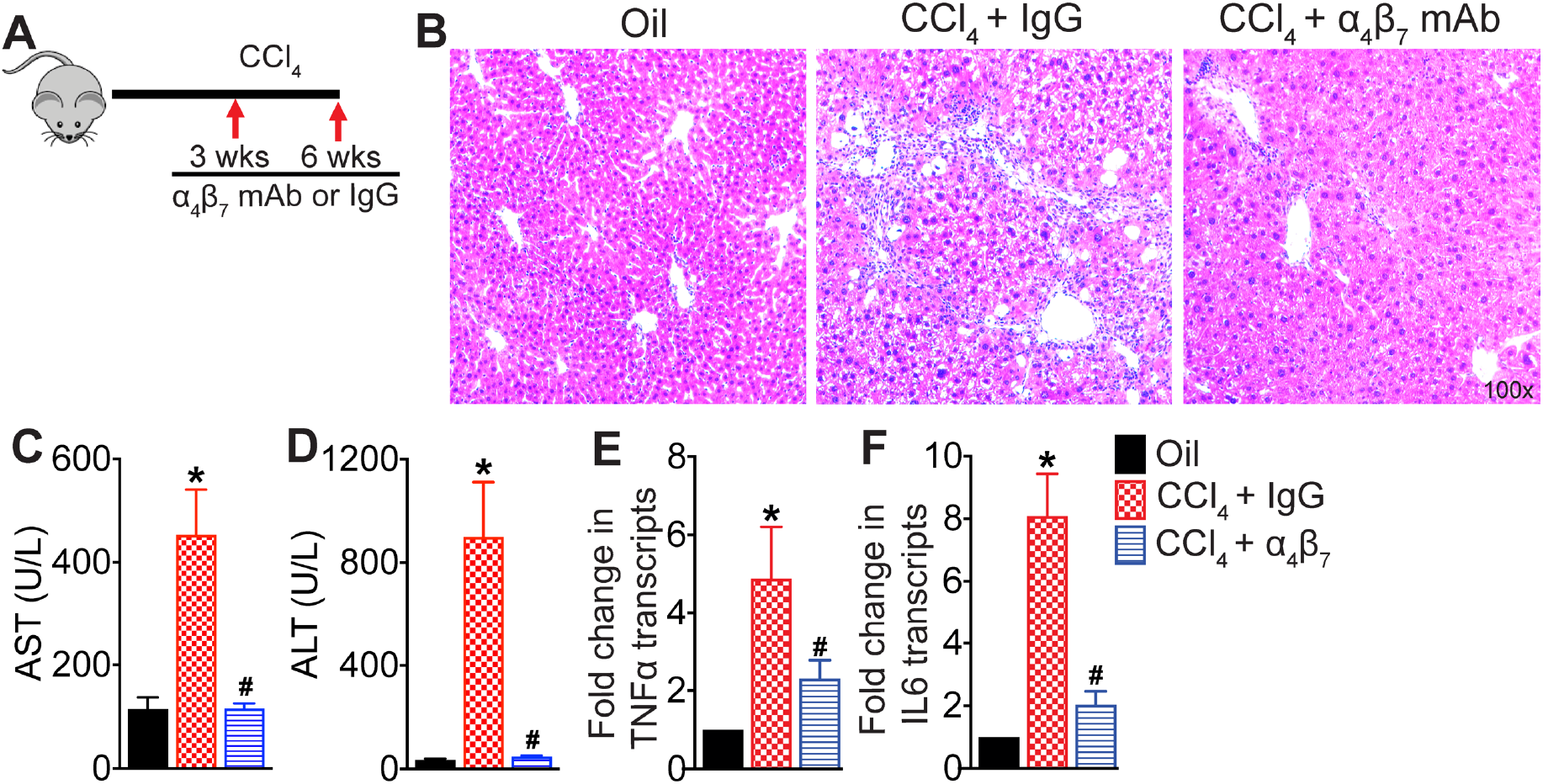
Integrin α_4_β_7_blockade attenuates CCl_4_-induced hepatic inflammation. (**A**) Schematic of study design. Liver fibrosis was induced by oral administration of CCl_4_ for six weeks. Mice treated with CCl_4_ for six weeks received α_4_β_7_ mAb or IgG isotype antibody for four weeks starting at week three. Mice administered the vehicle, olive oil, served as controls (n = 5 - 8 mice per group). (**B**) Representative photomicrographs of Hematoxylin and Eosin (H&E) stained liver tissue sections. (**C-D**) Serum AST and ALT levels. (**E-F**) Expression of key molecules associated with hepatic inflammation (n = 5 - 8 mice per cohort). Data are presented as mean ± SEM. Asterisks indicate significant differences (p < 0.05) between oil and IgG isotype control. Hashtags indicate significant differences (p < 0.05) between IgG and α_4_β_7_ mAb treatments.

### Integrin α_4_β_7_ blockade reduces hepatic fibrosis progression

To determine the effect of α_4_β_7_ blockade on hepatic fibrosis, liver tissue sections were stained with Sirius Red to assess collagen deposition while transcript levels of key molecules associated with hepatic fibrogenesis were quantified by qRT-PCR. As shown in **Fig. 2A**, CCl_4_ treatment induced severe hepatic fibrosis indicated by increased collagen deposition in the livers of CCl_4_ + IgG-treated mice compared to vehicle controls. The transcript levels of fibrogenesis marker α-smooth muscle actin (αSMA), a key marker of hepatic stellate cell (HSC) activation, collagen I (Col IA1 and Col IA2), transforming growth factor-β1 (TGF-β1), and tissue inhibitor of metalloproteinase-1 (TIMP-1) were also significantly elevated in the CCl_4_ + IgG-treated mice relative to controls (**Fig. 2B-F**). Treatment with α_4_β_7_ mAb reduced hepatic fibrosis reflected by decreased collagen deposition in the liver and significant reduction in the transcript levels of αSMA, Col IAI and Col IAII, TGF-β1 and TIMP-1 (**Fig. 2A-F**). These results were further corroborated by confocal imaging of αSMA-stained liver tissue sections demonstrating significant inhibition of HSC activation evidenced by reduced hepatic αSMA-expressing HSCs following α_4_β_7_mAb treatment (**Fig. 2G**). Taken together, these results demonstrate that α_4_β_7_ blockade attenuates CCl_4_-induced hepatic fibrosis.

**Figure 2.**
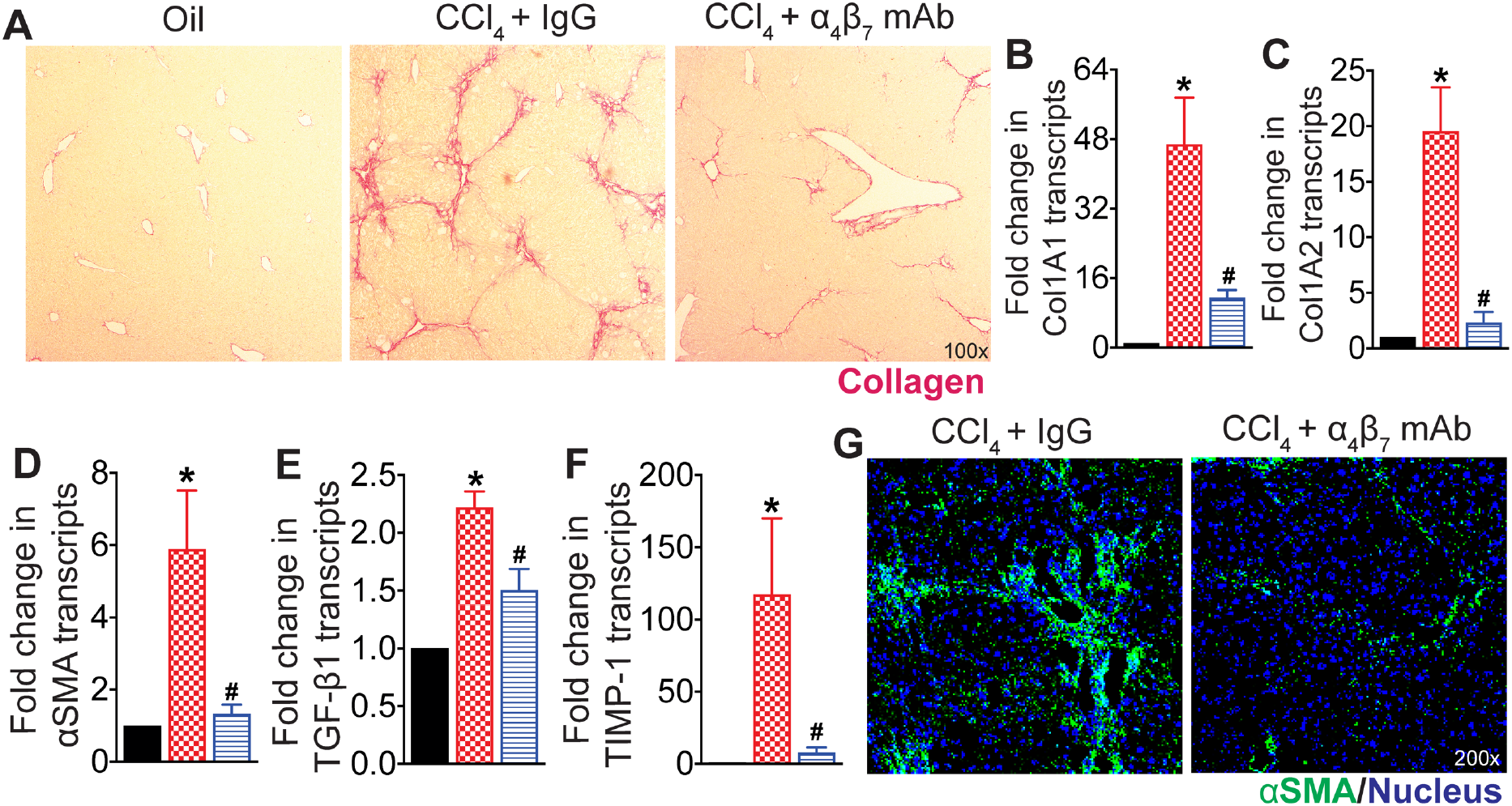
Integrin α_4_β_7_blockade attenuates CCl_4_-induced hepatic fibrosis progression. (**A**) Representative photomicrographs of Sirius Red-stained liver tissue sections. Liver fibrosis was induced by oral administration of CCl_4_ for six weeks. Mice treated with CCl_4_ for six weeks received α_4_β_7_ mAb or IgG isotype antibody for four weeks starting at week three. Mice administered the vehicle, olive oil, served as controls (n = 5 - 8 mice per group). (**B-F**) Expression of fibrosis markers in the liver. (**G**) Representative confocal images of αSMA (green) and DAPI (blue) immunofluorescence in the liver (n = 5 mice per group). Data are presented as mean ± SEM. Asterisks indicate significant differences (p < 0.05) between oil and IgG isotype control. Hashtags indicate significant differences (p < 0.05) between IgG and α_4_β_7_mAb treatments.

### α_4_β_7_ mAb treatment reduces accumulation of α_4_β_7_+ T cells in the fibrotic liver

To determine whether α4β7 mAb treatment reduces hepatic inflammation and fibrosis in CCl4-treated mice by decreasing the recruitment of α4β7+ immune cells to the liver, we assessed hepatic immune cell infiltrates using flow cytometry. Phenotypic analysis of intrahepatic lymphocytes revealed significant enrichment of α4β7+ CD4 and CD8 T cells in the livers of CCl4 + IgG-treated mice compared to vehicle controls (**Fig. 3B-E**). Treatment with α_4_β_7_ mAb significantly reduced intrahepatic α_4_β_7_+ CD4 and CD8 T cells in CCl_4_-treated mice compared to IgG controls (**Fig. 3B-E**). These findings were further supported by the significantly higher transcript levels of integrins α_4_, β_7_ and β_1_ in the CCl4 + IgG-treated mice relative to vehicle controls (**Fig. 3F-H**). Treatment with α_4_β_7_mAb significantly reduced transcript levels of integrins α_4_and β_7_, but not β_1_, demonstrating the specificity of α_4_β_7_mAb in blocking the recruitment of α_4_β_7_+ immune cells to the fibrotic liver (**Fig. 3F-H**). Colocalization studies also revealed increased infiltration of α_4_β_7_+ CD4 and CD8 T cells in the livers of CCl_4_ + IgG-treated mice relative to vehicle controls (**Fig. 3I-J**). Notably, treatment with α_4_β_7_ mAb markedly decreased the infiltration of α_4_β_7_+ CD4 and CD8 T cells in the liver relative to IgG-treated mice, demonstrating that α_4_β_7_ blockade reduces α_4_β_7_+ T cell recruitment to the fibrotic liver (**Fig. 3I-J)**. Collectively, these findings indicate that α_4_β_7_ regulates recruitment and accumulation of CD4 and CD8 T cells in the fibrotic liver.

**Figure 3.**
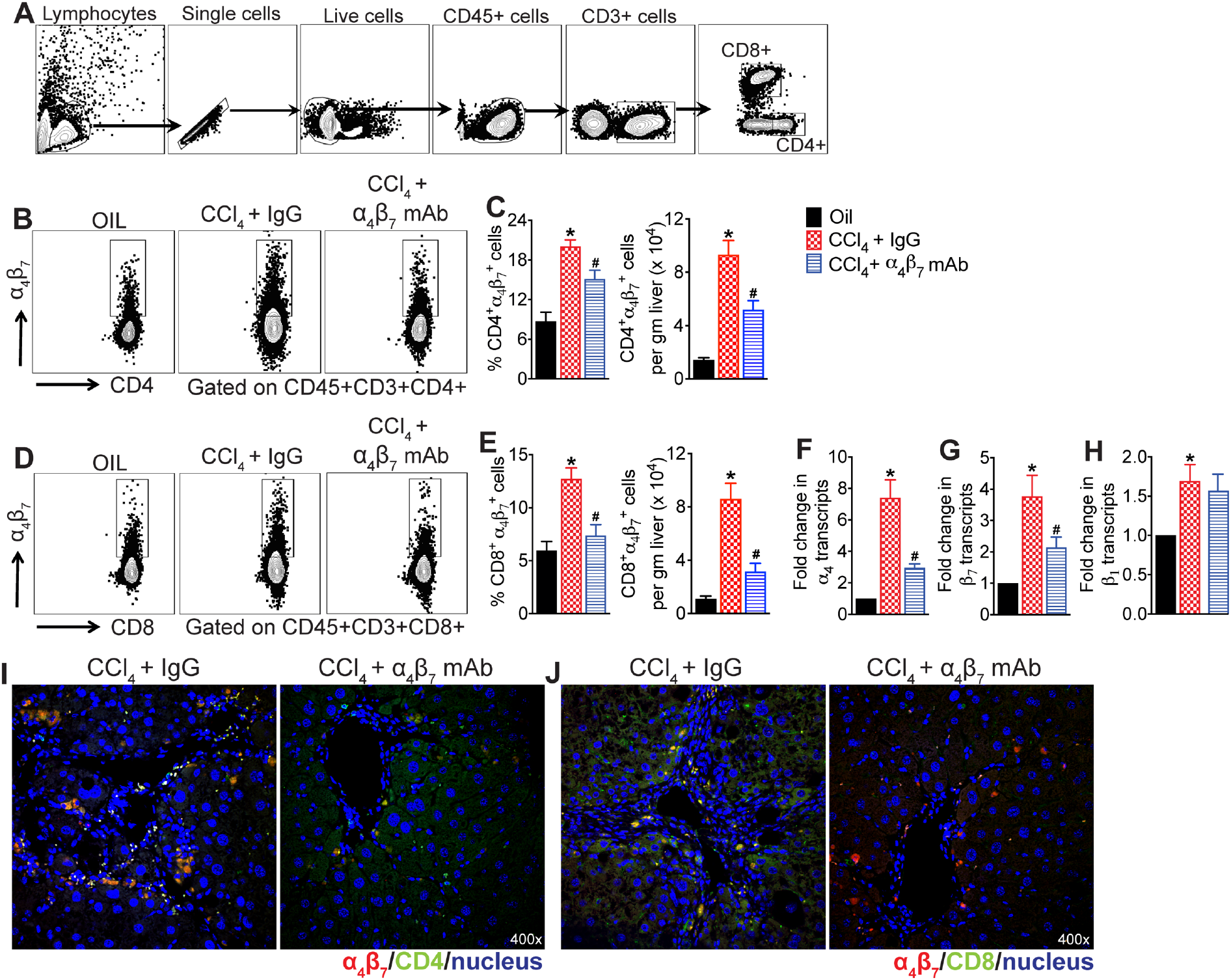
Integrin α_4_β_7_ mAb treatment reduces CCl_4_-induced infiltration of α_4_β_7_+ T cells in the liver. (**A**) Gating strategy used to identify intrahepatic lymphocytes. (**B-E**) Representative flow plots show percent of α_4_β_7_+ CD4 and CD8 T cells, and bar graphs show percentage and total number of α_4_β_7_+ CD4 and CD8 T cells in the liver. Mice treated with CCl_4_ for six weeks received α_4_β_7_ mAb or IgG isotype for four weeks starting at week three. Mice administered the vehicle, olive oil, served as controls (n = 5 - 8 mice per group). (**F-H**) Expression of integrins α_4_, *Itga4*, β_7,_ *Itgb7* and β_1,_ *Itgb1* in the liver. (**I-J**) Representative confocal images of (**I**) α_4_β_7_(red) and CD4 (green) and (**J**) α_4_β_7_(red) and CD8 (green) immunofluorescence in the liver. Nuclei are stained with DAPI (blue). Data are presented as mean ± SEM. Asterisks indicate significant differences (p < 0.05) between oil and CCl_4_ + IgG isotype controls. Hashtags indicate significant differences (p < 0.05) between IgG and α_4_β_7_ mAb treatments.

### MAdCAM-1 blockade reduces CCl_4_-induced hepatic infiltration of α_4_β_7_^+^ T cells

To determine if MAdCAM-1 blockade have a similar impact on hepatic α_4_β_7_+ T cell recruitment in CCl_4_-treated mice and ultimately attenuate liver injury, a cohort of the CCl_4_-treated mice were treated with MAdCAM-1 mAb for four weeks. Mice treated with IgG isotype served as controls (**Fig. 4A)**. As shown in **Fig. 4B-E**, four weeks of MAdCAM-1 mAb treatment significantly decreased intrahepatic α_4_β_7_+ CD4 and CD8 T cells in the CCl_4_ + MAdCAM-1 mAb-treated mice relative to IgG controls. Treatment with MAdCAM-1 mAb did not impact the frequency but reduced the total number of Foxp3+ T regulatory cells (Tregs) in the liver (**Fig. 4F**). Analysis of intrahepatic innate immune cells revealed significant reduction in the frequency and the total number of macrophages (CD3-CD11b+Ly6G-F4/80+ cells) and monocytes (CD3-CD11b+Ly6G-Ly6C+ cells), but not neutrophils (CD3-CD11b+Ly6G+ cells), in the liver of CCl_4_ + MAdCAM-1 mAb-treated mice relative to IgG controls (**Fig. 4G-I**). Treatment with MAdCAM-1 mAb significantly decreased the transcript levels of integrins α_4_and MAdCAM-1, but increased integrin β_7_transcripts, in the CCl_4_-treated mice relative to IgG controls (**Fig. 4J-L**). These findings demonstrate that, similar to α_4_β_7_, MAdCAM-1 blockade also reduces the recruitment of α_4_β_7_+ T cells to the fibrotic liver, suggesting a role for the α_4_β_7_/MAdCAM-1 axis in regulating α_4_β_7_+ T cells to the fibrotic live.

**Figure. 4.**
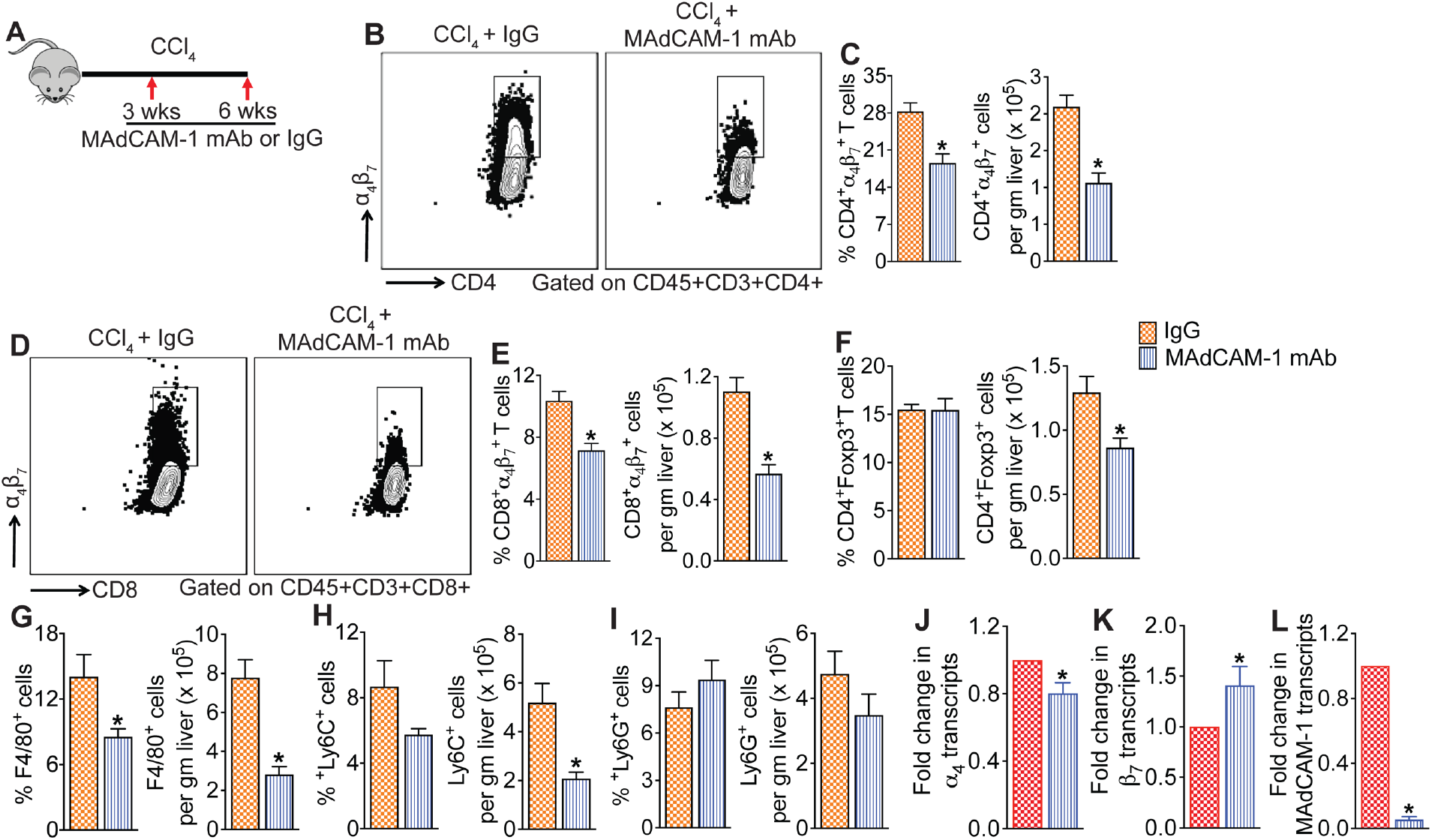
MAdCAM-1 blockade reduces CCL_4_-induced infiltration of α_4_β_7_+ T cells in the liver. (**A**) Schematic of study design. Mice treated with CCl_4_ for six weeks received MAdCAM-1 mAb or IgG isotype antibody for four weeks starting at week three (n = 5 - 8 mice per cohort). (**B-E**) Representative flow plots show percent of α_4_β_7_+ CD4 and CD8 T cells, and bar graphs show percentage and total number of α_4_β_7_+ CD4 and CD8 T cells in the liver. (**F**) Percentage and total number of Foxp3+ CD4 T cells in the liver. (**G-I**) Percentage and total number of macrophages (CD3-CD11b+Ly6G-F4/80+ cells), monocytes (CD3-CD11b+Ly6G-Ly6C+ cells) and neutrophils (CD3-CD11b+Ly6G+ cells) in the liver. (**J-L**) Expression of integrins α_4_, *Itga4* and β_7,_ *Itga7* and *Madcam-1* in the liver (n = 5). Data are presented as mean ± SEM. Asterisks indicate significant differences (p < 0.05) between IgG and MAdCAM-1 mAb treatments.

### α_4_β_7_+ CD4 T cells enriched for markers of activation and proliferation demonstrating an effector phenotype

To determine the function of α_4_β_7_+ CD4 T cells, we assessed the expression of T cell activation and proliferation markers in α_4_β_7_ + and α_4_β_7_ - CD4 T cell subsets in the liver of CCl_4_ + IgG treated mice. Our analysis revealed that a significantly higher frequency of α_4_β_7_+ CD4 T cells expressed the T cell activation marker CD44, acute T cells activation marker CD69, the Th1 transcriptional factor driving IFN production Tbet, and the cell proliferation marker Ki67 compared to the α_4_β_7_-CD4 T cells (**Fig 5A-D**). These data suggest that α_4_β_7_+ CD4 T cells comprise effector T cells with capacity for proliferation and cytokine production.

**Figure. 5.**
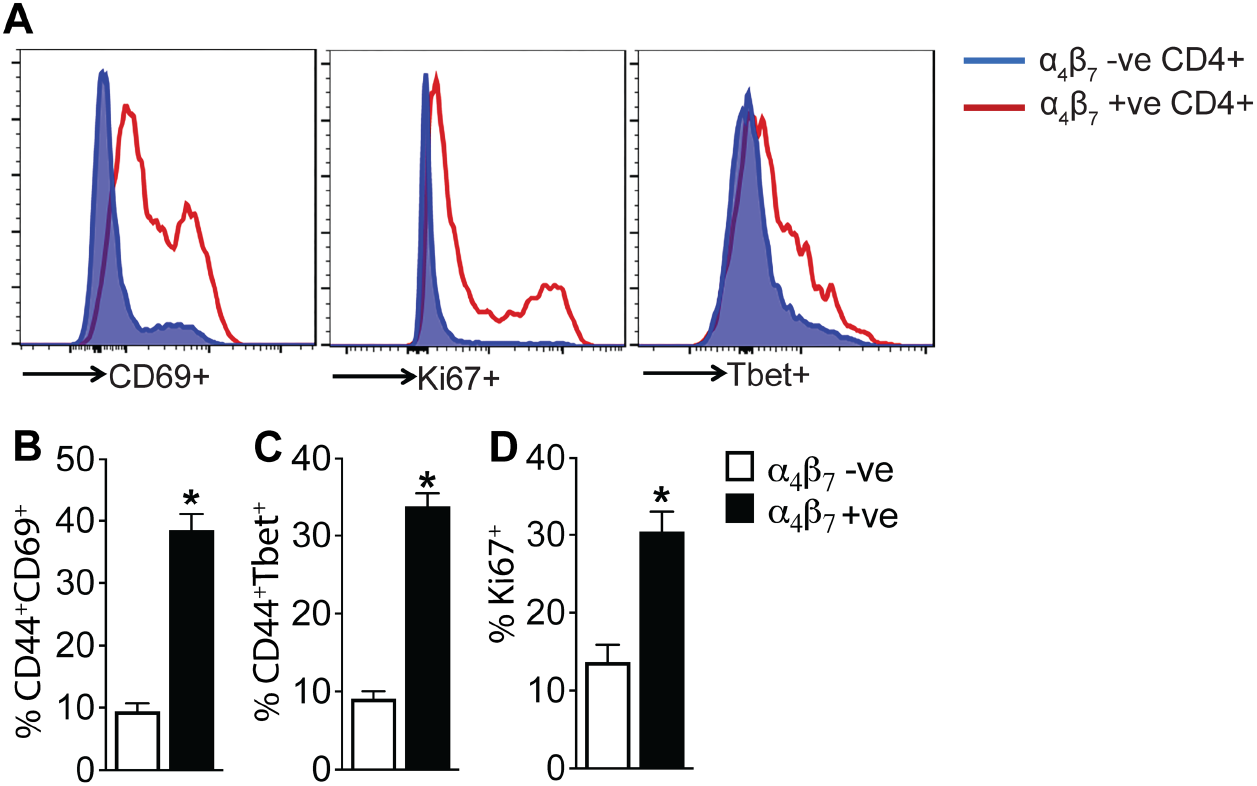
α_4_β_7_+ CD4 T cells enriched for markers of activation and proliferation. demonstrating an effector phenotype. (**A**) Representative histograms show median fluorescence intensity of stated markers in the liver of mice treated with CCl_4_ for six weeks. Gated on CD45+CD3+CD4+ cells. Mice were treated with IgG during the final four weeks (n = 5). Bar graphs show percentage of α_4_β_7_+ and α_4_β_7_-CD4 T cells expressing (**B**) T cell activation markers CD44 and CD69, (**C**) Th1 cell marker Tbet and (**D**) proliferation marker Ki67 in the liver. Data are presented as mean ± SEM. Asterisks indicate significant differences (p < 0.05) between α_4_β_7_+ and α_4_β_7_-CD4 T cells.

### MAdCAM-1 blockade attenuates CCl_4_-induced hepatic injury and fibrosis

Given the inhibitory effect of MAdCAM-1 blockade on the CCl_4_-induced recruitment of immune cells to the liver, we examined whether MAdCAM-1 blockade reduced CCl_4_-induced liver injury. Histological analysis of H&E-stained liver tissue sections revealed a notable decrease in hepatic inflammation and injury in CCl_4_ + MAdCAM-1-mAb treated mice relative to the CCl_4_ + IgG-treated mice (**Fig. 6A**). Improvement in hepatic injury in CCl_4_ + MAdCAM-1-mAb treated mice was confirmed by decreased serum ALT levels (**Fig. 6B**). Treatment with MAdCAM-1-mAb significantly decreased the transcript levels of TGF-β and IL6, but not TNF-α (**Fig. 6C-E**). Treatment with MAdCAM-1 mAb decreased hepatic fibrosis indicated by marked decrease in the deposition of hepatic collagen in the Sirius Red-stained liver tissue sections (**Fig. 6F**). Transcript levels of key molecules associated with fibrogenesis of the liver, αSMA, TIMP-1, and Col IAI and Col IAII were significantly lower in the MAdCAM-1-mAb-treated mice relative to IgG controls (**Fig. 6G-J**). These data demonstrate that MAdCAM-1 blockade attenuates CCl_4_-induced hepatic inflammation and fibrosis.

**Figure. 6.**
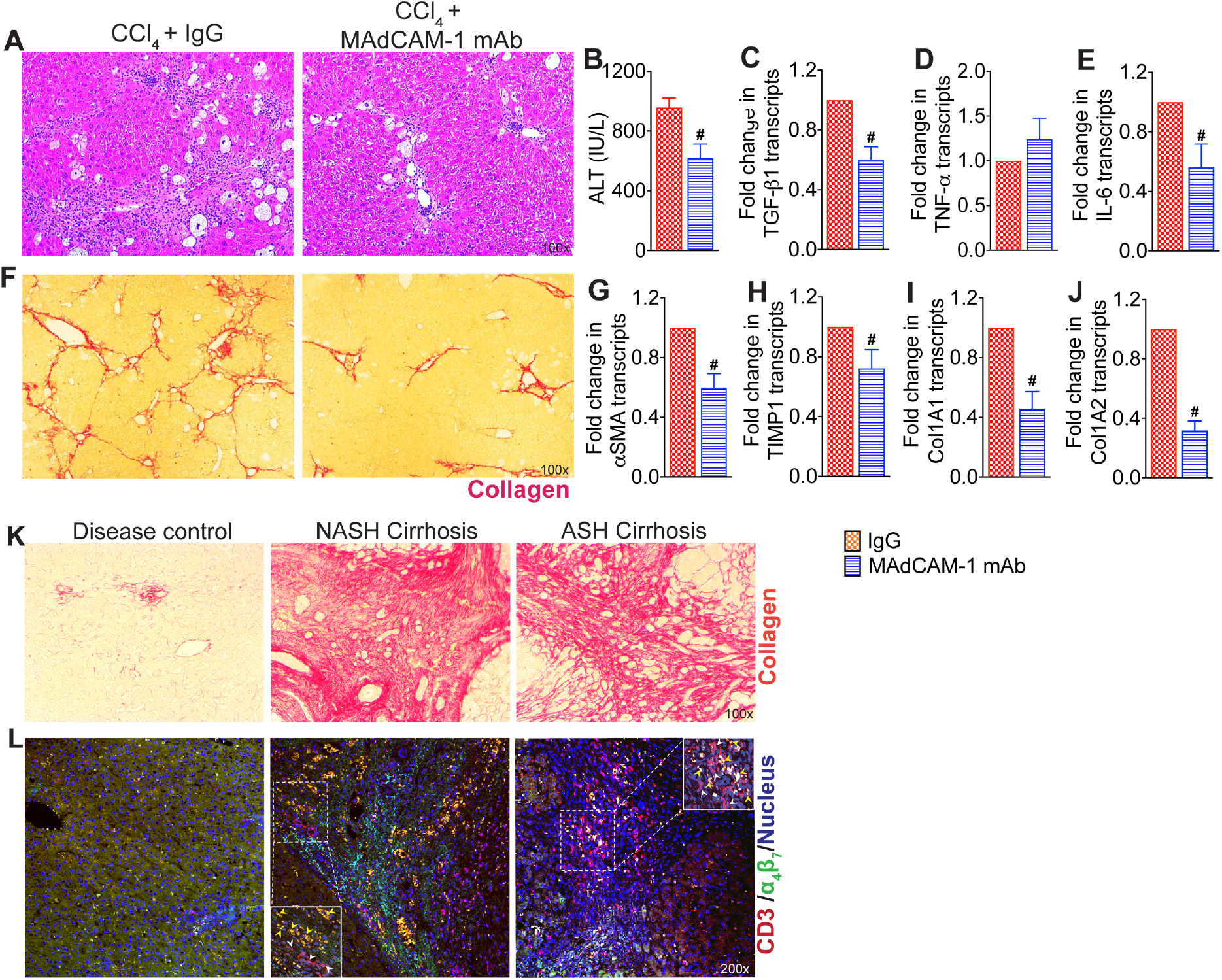
MAdCAM-1 blockade attenuates hepatic inflammation and fibrosis progression. (**A**) Representative photomicrographs of Hematoxylin and Eosin (H&E) stained liver tissue sections. Mice treated with CCl_4_ for six weeks received MAdCAM-1 mAb or IgG isotype antibody for four weeks starting at week three (n = 5 - 8 mice per group). (**B**) Serum ALT level. (**C-E**) Expression of key molecules associated with hepatic inflammation (n = 5-8 mice per group). (**F**) Representative photomicrographs of Sirius Red-stained liver tissue sections. (**G-J**) Expression of key fibrosis markers in the liver. Data are presented as mean ± SEM. Hashtags indicate significant differences (p < 0.05) between IgG and MAdCAM-1 mAb treatments. (**K**) Representative photomicrographs of Sirius Red-stained liver liver tissue sections from disease controls and people with nonalcoholic steatohepatitis (NASH) and alcoholic steatohepatitis (ASH)-associated cirrhosis. (**L**) Representative confocal images of CD3 (red), α4β7 (green) and DAPI (blue) immunofluorescence in the liver tissue sections. Insets show zoomed-in view of the region in white dotted box. White arrowheads, CD3 T cells. Yellow arrowheads, α_4_β_7_+ CD3 T cells.

### Increased infiltration of α_4_β_7_+ T cells in the livers of NASH- and ASH-associated cirrhosis patients

To determine whether α_4_β_7_+ T cells are also involved in the hepatic inflammation and fibrosis in people with end-stage liver disease, we probed liver tissue sections obtained from 10 people with NASH-associated cirrhosis and 10 people with ASH-associated cirrhosis for the presence of α_4_β_7_+ T cells using confocal laser scanning microscopy. Liver tissue from tumor-adjacent parenchyma was used as controls. Staining of liver tissue sections with α_4_β_7_ and CD3 antibodies revealed higher infiltration of α_4_β_7_ and CD3 double-positive T cells and the formation of large inflammatory aggregates of α_4_β_7_+ T cells in the livers of people with NASH- and ASH-associated cirrhosis (**Fig. 6K-L**). Interestingly, higher inflammatory aggregates of α_4_β_7_+CD3+ T cells were observed in the fibrotic septa of livers from people with NASH-associated cirrhosis compared to those with ASH-associated cirrhosis. Collectively, these findings suggest the involvement of α_4_β_7_+ T cells in people with end-stage liver disease.

## DISCUSSION

Immune cells recruited to the injured liver are the primary instigators of sustained chronic inflammation and resulting fibrosis ^12–15^; yet we lack complete understanding of the molecular mechanisms involved in immune-cell recruitment and the role of various immune cell subsets in promoting hepatic fibrosis. In this study, we report the involvement of the α_4_β_7_/MAdCAM-1 axis in the recruitment of α_4_β_7_+ T cells to the fibrotic liver, highlighting their pivotal role in promoting hepatic fibrosis progression. Using an established mouse model of liver fibrosis, we demonstrate the enrichment of α_4_β_7_+ T cells in the fibrotic liver, and importantly, show that blocking α_4_β_7_ or MAdCAM-1 reverses fibrosis progression by reducing the recruitment of α_4_β_7_+ T cells to the liver. These findings unveil a previously unappreciated role for α_4_β_7_+ T cells in promoting hepatic fibrosis progression.

Our clinical findings revealing the presence of inflammatory aggregates of α_4_β_7_+ T cells in the livers of individuals with cirrhosis strongly indicate their involvement in liver injury. Furthermore, our data indicating larger inflammatory aggregates of α_4_β_7_+ T cells in NASH-associated cirrhosis compared to ASH-associated cirrhosis suggest a potential heightened role of these cells in contributing to liver injury in NASH-associated cirrhosis. These findings, combined with previous reports on the α_4_β_7_/MAdCAM-1 axis in regulating T cell recruitment in various CLDs, underscore the critical contribution of α_4_β_7_+ T cells to hepatic inflammation and fibrosis progression in CLD ^8, 10, 16, 17^.

Our findings from an established mouse model of liver fibrosis recapitulated our clinical findings by demonstrating that α_4_β_7_+ T cells are actively recruited to the fibrotic liver and α_4_β_7_+ T cells contribute to driving hepatic fibrosis progression in this model. Our results align with previous studies that reported protective effects against diet-induced non-alcoholic steatohepatitis (NASH) in mice lacking whole-body expression of MAdCAM-1 or treated with α_4_β_7_or MAdCAM-1 mAbs ^6, 18^. Additionally, whole-body knockout mice lacking MAdCAM-1 or β_7_ have been shown to be protected from concavalin A-induced hepatitis ^19^. In contrast, whole-body β_7_knockout mice have been shown to exhibit enhanced susceptibility to diet-induced NASH ^18^. These discrepancies may be attributed to inherent defects in the development of gut-associated lymphoid tissue in β_7_ knockout mice, making data interpretation more complex ^20, 21^. These conflicting findings also emphasize the limitations of relying solely on whole-body knockout mouse models in functional research, as non-specific phenotypes can often be overlooked. It should be noted that the α_4_β_7_mAb we used for our studies is a highly specific antibody that specifically binds to a conformational epitope accessible only in the α_4_β_7_ heterodimer ^22–24^. The specificity of this antibody has been extensively evaluated across various experimental and clinical contexts, and the humanized version of the α_4_β_7_ monoclonal antibody has been recognized as an effective treatment for inflammatory bowel disease ^23–26^. Furthermore, we demonstrate that treatment with MAdCAM-1 mAb also protects mice from CCl_4_-induced liver fibrosis by blocking α_4_β_7_+ T cell recruitment to the liver. These findings not only underscore the significance of the α_4_β_7_/MAdCAM-1 axis in modulating T cell recruitment to the liver but also highlight the critical role played by α_4_β_7_+ T cells in promoting hepatic fibrosis.

Our findings that α_4_β_7_/MAdCAM-1 axis regulate CD4 T cell recruitment to the fibrotic liver adds to our previously reported role of this axis in regulating CD4 T cell recruitment to the NASH liver ^6^. Consistent with earlier reports, our results confirmed that a significant proportion of α_4_β_7_+ CD4 T cells are effector T cells ^27, 28^. Effector CD4 T cells are known to contribute to hepatic inflammation and fibrosis through the secretion of proinflammatory and profibrotic cytokines. Their activation leads to the stimulation of both innate and adaptive immune cells, thereby perpetuating the inflammatory and fibrotic processes within the liver ^29–31^. In contrast, regulatory T cells expressing the transcription factor Foxp3 have been recognized for their ability to suppress immune responses and promote tissue repair ^32^. In our study, we found that the α_4_β_7_/MAdCAM-1 axis specifically targets the recruitment of α_4_β_7_+ effector CD4 T cells to the fibrotic liver, while sparing the migration of Foxp3+ Tregs. This selectivity in blocking effector CD4 T cell recruitment while preserving the presence of immunomodulatory Tregs may have played a crucial role in the effectiveness of α_4_β_7_and MAdCAM-1 mAbs in suppressing hepatic fibrosis.

The involvement of CD8 T cells in tissue injury and fibrosis is of particular importance as antigen-specific or bystander activation of CD8 T cells can promote tissue injury by cytolysis of damaged cells or exacerbate inflammation by producing proinflammatory cytokines including TNF-α, INF-γ, IL-13, and IL-4 ^33–37^, which promote recruitment of proinflammatory immune cells to the injured tissue. Our data provide evidence supporting the role of the α_4_β_7_/MAdCAM-1 axis in regulating the recruitment of CD8 T cells to the liver. Specifically, we demonstrate that a significant subset of CD8 T cells in the fibrotic liver express α_4_β_7_. We demonstrate that blocking antibodies against α_4_β_7_ and MAdCAM-1 that reduce recruitment of α_4_β_7_+ CD8 T cells to the liver improves CCL_4_-induced hepatic fibrosis. These findings expand upon the previously reported roles of the α_4_β_7_/MAdCAM-1 axis in regulating CD4 T cell homing to the liver in NASH ^6^ and PSC ^17^. Furthermore, our findings suggest that α_4_β_7_-expressing CD8 T cells play a prominent role in promoting hepatic fibrosis in CLD. However, considering previous reports that CD8 T cells play a protective role during hepatic injury resolution by eliminating activated HSCs ^38^, and their dysfunction at the advanced stages of CLD exacerbate disease progression ^39, 40^, further comprehensive investigations are needed to understand the specific functions of different CD8 T cell subsets in promoting and regressing hepatic fibrosis in CLD.

In conclusion, our findings shed light on the crucial role of the α_4_β_7_/MAdCAM-1 axis in in regulating hepatic fibrosis and highlight the potential therapeutic strategy of utilizing mAbs targeting integrin α_4_β_7_ or MAdCAM-1 to treat inflammation and fibrosis in CLD.

### Data availability statement

Data sharing not applicable to this article as no datasets were generated or analyzed during the current study.

### Author Contributions

Conceptualization: RR; Methodology: RR, SSI, BG, RPR, PBP, SC, AC, SS, ADS; Investigation: BG, SSI, RPR, PBP; Visualization: RR, BG, RPR, SSI, PBP; Funding acquisition: RR; Project administration: RR, BG, RPR; Supervision: RR; Writing – original draft: BG, RR; Writing – review & editing: RR, SSI, ADS, SPM.

## Acknowledgements

Research reported in this publication was supported by the National Institute of Diabetes and Digestive and Kidney Diseases of the National Institutes of Health under award number K01DK110264 and R01DK124351 to RR; R01DK62277, R01DK100287 and Endowed Chair for Experimental Pathology to SPM; and R56AI150409 and RF1AG06001 to SSI. This research project was also supported in part by the University of Pittsburgh Clinical Biospecimen Processing and Repository Core and Advanced Cell and Tissue imaging Centre of the Pittsburgh Liver Research Centre supported by NIH/NIDDK Digestive Disease Research Core Center grant P30DK120531 and the University of Pittsburgh Unified Flow Core through the resources provided.

